# The Extracellular Matrix Limits Nanoparticle Diffusion and Cellular Uptake in a Tissue-Specific Manner

**DOI:** 10.1101/2024.07.30.605895

**Authors:** Devorah Cahn, Alexa Stern, Michael Buckenmeyer, Matthew Wolf, Gregg A. Duncan

## Abstract

Overexpression and remodeling of the extracellular matrix (ECM) in cancer and other diseases may significantly reduce the ability of nanoparticles to reach target sites, preventing effective delivery of therapeutic cargo. Here, we evaluate how tissue-specific properties of the ECM affect nanoparticle diffusion using fluorescence video microscopy and cellular uptake via flow cytometry. In addition, we determined how PEGylation chain size and branching influence the ability of nanoparticles to overcome the ECM barrier from different tissues. We found that purified collagen, in the absence of other ECM proteins and polysaccharides, presented a greater barrier to nanoparticle diffusion as compared to decellularized ECM from the liver, lung, and small intestine submucosa. Nanoparticles with dense PEG coatings achieved up to ∼2000-fold enhancements in diffusion rate and cellular uptake up to ∼5-fold greater than non-PEGylated nanoparticles in the presence of ECM. We also found nanoparticle mobility in the ECM varied significantly between tissue types and the optimal nanoparticle PEGylation strategy to enhance ECM penetration was strongly dependent on ECM concentration. Overall, our data supports the use of low molecular weight PEG coatings which provides an optimal balance of nanoparticle penetration through ECM and uptake in target cells. However, tissue-specific enhancements in ECM penetration and cellular uptake were observed for nanoparticles bearing a branched PEG coating. These studies provide new insight into tissue specific ECM barrier functions which can facilitate the design of nanoparticles that effectively transport through target tissues, improving their therapeutic efficacy.

## Introduction

Nanoparticle (NP) drug carriers have proven to be an effective tool in increasing the bioavailability and efficacy of drugs by preventing drug degradation and facilitating entry into target cells.^1^ However, extracellular barriers have proven to be significant obstacles to the transport of these carriers, reducing their overall efficacy. One such barrier is the extracellular matrix (ECM) which often plays an important role in the progression and resolution of many diseases.^2^ The ECM is a structural and protective matrix that surrounds cells and is composed of collagens, proteoglycans, fibronectin, elastin, laminins, and other glycoproteins.^1,3,4^ NPs must diffuse through this network to reach their target cells and deliver their cargo.^5^ The network can obstruct NP transport via steric and electrostatic interactions which results in NP entrapment within the mesh and altered therapeutic biodistribution.^5–8^

Many tissues undergo ECM remodeling during disease, which can further hinder the transport of NPs through the matrix.^9,10^ One important example of this is the upregulation of ECM components in many cancers.^2,11,12^ Both the primary tumor site and distant metastatic niche undergo significant ECM changes during progression.^13–15^ Studies indicate that fewer than 1% of NPs reach target cancer cells with the majority accumulating at the tumor periphery.^16^ The ECM plays a key role in cancer progression and its resistance to immune cell and drug targeting.^17^ Moreover, it has been shown the genes for ECM proteins are upregulated in many drug-resistant cancers.^18^ The overexpression of the various components in the ECM within the tumor microenvironment increases the stiffness of the matrix and reduces its pore size.^19^ This serves to further trap NPs within the matrix via steric hindrance, preventing them from reaching their target.^5,6,20^ For example, hyaluronic acid (HA) is a glycosaminoglycan (GAG) that is commonly overexpressed by cancer and stromal cells.^2^ Upregulation of charged ECM components such as HA can reduce NP diffusivity via electrostatic interactions and by inducing an increase in interstitial fluid pressure.^21^ Polyethylene glycol (PEG) is often used to modify the surfaces of NPs to aid in NP transport through the ECM and other hydrogel matrices.^22^ Studies have shown that PEGylation density, molecular weight (MW), and architecture can affect NP transport through models of the ECM where increased density, lower MW, and PEG branching improved NP mobility.^7,20,22,23^ However, these studies generally focused on collagen and Matrigel models which are not wholly representative of the native matrix.

The ECM can be subdivided into the basement membrane, a dense layer of ECM which lies directly on top of epithelial and endothelial cell membranes, and the interstitial matrix surrounding it.^24,25^ These two elements of the ECM possess different compositions and varieties of collagens and proteoglycans.^4,25^ They also have different mechanical properties (e.g. stiffness) and density that are important factors in NP transport.^19^ As a basement membrane model, Matrigel is primarily composed of laminin as well as collagen IV with a stiffer and lower porosity network compared to the interstitial matrix.^19,26,27^ The interstitial matrix on the other hand is primarily composed of fibrillar collagens including Type I, III, and V and has a more moderate porosity and lower stiffness.^19,25^ However, the types of collagens and their organization within the interstitial matrix varies depending on the type of tissue where they are found.^28^ It is currently unclear to what extent these tissue-specific characteristics affect NP transport through ECM and how NP PEGylation impacts ECM penetration in distinct tissues.

In this work, we investigated whether tissue specific properties of ECM affected NP transport and whether the optimal PEGylation strategy is affected by these tissue-specific differences. To accomplish this goal, we developed a hydrogel model of the interstitial matrix by utilizing decellularized ECM (dECM) from three different tissues: lung, liver, and small intestine submucosa (SIS). This system isolates tissue-specific ECM composition and enables control over ECM concentration to model changing matrix density. Lung and liver ECM were chosen for these studies as these tissues are common sites of nanoparticle accumulation following systemic administration.^29–31^ The lung and liver are also often sites of cancer metastasis and other fibrotic diseases associated with significant ECM remodeling^32–35^ which may effectively limit NP-based drug delivery systems. SIS has been used extensively in the field of tissue engineering, providing a basis for comparison to the other tissue types.^36,37^ We evaluated the transport and cellular uptake of NPs modified with either linear or branched PEG in these hydrogels to determine optimal PEGylation strategies depending on the characteristics of the ECM within target tissues.

## Materials and Methods

### Materials

Tris, ethylenediaminetetraacetic acid (EDTA), sodium deoxycholate (SDC), Triton X-100, and peracetic acid (PAA) were obtained from Sigma Aldrich. Trypsin (Gibco) was obtained from ThermoFisher. Fluorescent carboxylate-modified polystyrene (PS-COOH) nanoparticles were obtained from Thermo Scientific. 2 kDa and 5 kDa linear methoxy PEG-amine were purchased from Creative PEGWorks. 10 kDa linear methoxy PEG-amine and 10 kDa 4-arm PEG amine were obtained from JenKem Technologies.

### Lung and Liver Decellularization

All tissues used in this study were sourced from market weight pigs (Tissue Source LLC, Zionsville, IN) in compliance with ISO 13485. dECM was obtained from porcine lung and liver through a series of chemical and mechanical treatments. Frozen tissue was thawed and diced into pieces smaller than 0.5 mm after which it was placed in a hypotonic solution consisting of 10 mM Tris and 5 mM EDTA for 30 minutes. The solution was then replaced with a hypertonic solution consisting of 50 mM Tris, 1 M NaCl, and 10 mM EDTA. After 30 minutes, the solution was replaced with a 0.02% trypsin with 0.05% EDTA solution for one hour. The tissue was washed in water, PBS, and then water again and then 4% SDC was added to the tissue and agitated in the solution for 3 hours. The tissue was then washed and subjected to 3% Triton X-100 for 3 hours. Finally, 0.1% PAA was added to the tissue and the resulting dECM was frozen, lyophilized, and cryomilled.

### Small Intestine Submucosa Decellularization

Decellularized small intestine submucosa was prepared as previously reported.^38^ Briefly, gastrointestinal fluids were flushed from porcine small intestines and cut open lengthwise. The mucosal and muscularis layers were then removed by mechanically delaminating the tissue. The submucosa was cut into 6 inch pieces and further decellularized and sterilized in a solution of 4% alcohol and 0.1% PAA. The decellularized tissue was then washed using 1x PBS and Type 1 water after which it was lyophilized and cryomilled in preparation for hydrogel formulation.

### dECM Hydrogel Formation

To create dECM hydrogels, cryomilled dECM was digested in 1 mg/mL pepsin for 48 hours at an ECM concentration of 10 mg/mL and kept cold as previously described.^39^ The digested dECM was neutralized using NaOH and brought to isotonic salt concentrations with 10X PBS to form a pre-gel that was diluted concentrations of 4, 6, or 8 mg/mL using PBS for multiple particle tracking studies or media for cellular uptake studies. The pre-gel was then incubated at 37°C for at least 30 minutes to form the final dECM hydrogel.

### Biochemical characterization of ECM components

The soluble collagen content of the decellularized tissue was determined using a Sircol Soluble Collagen Assay kit (Biocolor S1000). The dye was prepared by dissolving Sirius red F3B in a saturated aqueous solution of Picric acid at a concentration of 1 mg/mL at pH 2-3. Nutragen, a type I bovine collagen, was diluted to different concentrations to use as a standard curve. Pepsin digested dECM was used to isolate soluble collagen. 100 uL of either the collagen standard or dECM was mixed with 1 mL of dye for 30 minutes at 4°C. The mixture was then centrifuged, and the pellet was resuspended in 750 uL of ice-cold 0.1 M HCl. The solution was then centrifuged, and the pellet was resuspended in 1 mL of 0.5 M NaOH. The standards and samples were then added to a 96-well plate and the absorbance was read at 556 nm.

Sulfated glycosaminoglycan (sGAG) was also characterized in dECM from each tissue source. A papain digestion solution was prepared as a phosphate buffer consisting of 0.1 M disodium phosphate and 0.01 M disodium EDTA at a pH of 6.5. The buffer was used to create a 0.01 M L-cysteine solution which was combined with 25 mg/mL papain. Tissue was digested in the papain solution at a concentration of 10 mg/mL for 12-16 hours at 60°C. The digested tissue was added to a 96-well plate along with a standard curve of chondroitin sulfate with concentrations ranging from 0 to 125 μg/mL. The DMMB dye solution was prepared by combining 3.2 mg/mL DMMB in ethanol with 0.05 M NaCl, and 0.05 M glycine. The dye solution was diluted with water to a final DMMB concentration of 0.02 mg/mL. 250 μL of dye was added to 50 μL of sample in each well and the absorbance was read at a wavelength of 525 nm.

### Bulk Rheology of Hydrogels

The viscoelastic properties of the dECM hydrogels were obtained through dynamic rheological measurements using an ARES G2 rheometer (TA Instruments) with a 40-mm-diameter parallel plate geometry at 37°C. A strain sweep measurement from 0.02 to 100% strain at a frequency of 1 rad s^-1^ was used to determine the viscoelastic region of the hydrogels. The elastic modulus, G’ (ω), and viscous modulus, G” (ω) were collected using a frequency sweep measurement at a 0.5% strain and angular frequencies from 0.02 to 200 rad s^-1^. A time sweep measurement was conducted over 40 minutes at 0.5% strain and 0.2 rad s to determine differences in the gelation process between the hydrogels. Three replicate samples were analyzed for each type of hydrogel.

### Microrheology of hydrogels

Carboxylate-modified 500 nm fluorescent polystyrene (PS) nanoparticles (PS-COOH) were coated with 5 kDa PEG via a carboxyl-amine linkage as described in previous work.^23^ Size and zeta potential was measured in 10 mM NaCl at pH 7 using a Nanobrook Omni (Brookhaven Instruments). Nanoparticle diffusion in the dECM hydrogels was evaluated as described in our previous work.^23,40^ Briefly, 25 μL of either lung, liver, or SIS pre-gel was mixed with 1 μL of 0.002% w/v nanoparticles and allowed to gel at 37°C for 30 minutes. The diffusion of NPs was then imaged in real-time using a Zeiss 800 LSM microscope with a 63× water-immersion objective and Zeiss Axiocam 702 camera at a frame rate of 33.33 Hz for 10 seconds. Three independent samples were analyzed for each type of hydrogel and at least 3 high-speed videos were recorded for each sample. NP trajectories and diffusion rate were determined using a MATLAB-based image analysis software to track NP position over time. The time-averaged mean squared displacement (MSD; ⟨Δ*r*^2^(τ)⟩) as a function of lag time, τ, was calculated from these trajectories as ⟨Δ*r*^2^(τ)⟩ = ⟨[*x*(*t* + τ) − *x*(*t*)]^2^ + [*y*(*t* + τ) − *y*(*t*)]^2^⟩. The generalized Stokes-Einstein relation was then used to determine the viscoelastic properties of the hydrogels as described previously.^41^

### Multiple particle tracking (MPT)

Carboxylate-modified 100 nm fluorescent polystyrene (PS) nanoparticles (PS-COOH) were coated with either 2 kDa, 5 kDa, or 10 kDa linear PEG or with 10 kDa 4-arm branched PEG via a carboxyl-amine linkage as described in previous work. ^23,40^ Zeta potential was measured in 10 mM NaCl at pH 7 using a Nanobrook Omni (Brookhaven Instruments). Nanoparticle diffusion in the dECM and collagen hydrogels was evaluated as described in our previous work. Briefly, 25 μL of either lung, liver, SIS, or collagen pre-gel was mixed with 1 μL of 0.002% w/v nanoparticles and allowed to gel at 37°C for 30 minutes. The diffusion of NPs was imaged in real-time using a Zeiss 800 LSM microscope with a 63× water-immersion objective and Zeiss Axiocam 702 camera at a frame rate of 33.33 Hz for 10 seconds. NP trajectories and diffusion rate as measured by MSD were determined using a MATLAB-based image analysis software as described above.

### Nanoparticle Cellular Uptake

Human lung epithelial A549 cells were seeded on plastic at ∼3000 cells/cm^2^ in Ham’s F-12K (Kaighn’s) medium and incubated at 37°C and 5% CO_2_. Cells were passaged upon reaching 70-80% confluency at which time cells were dissociated from the plate using 0.05% trypsin EDTA for five minutes at 37 °C. To quantify nanoparticle uptake in A549 cells in 2D culture, cells were seeded in 24-well plates at 18,750 -cells/cm^2^ and allowed to adhere to the plate surface overnight. The medium in each well was then replaced with 0.5 mL of medium containing nanoparticles (∼10^11^ NPs/mL). The cells were then incubated with the nanoparticles for 4 hours after which the media was removed, and the cells were washed with phosphate-buffered saline (PBS) 3 times to remove excess nanoparticles. The cells were dissociated from the wells using 0.05% trypsin EDTA, collected via centrifugation, and then stained with Zombie NIR dye at a concentration of 1:1000 for 25 minutes on ice. The cells were then washed and fixed using 2% PFA on ice for 20 minutes after which they were washed again and stored in PBS for flow cytometry. To quantify nanoparticle uptake in A549 cells in 3D culture, cells were mixed with 4, 6, or 8 mg/mL lung, liver, or SIS pre-gels. 0.25 mL of the cell-gel mixture were added per well in 24-well plates. These cells were then incubated at 37°C for a minimum of 30 minutes to allow for gel formation after which 0.25 mL of media was added on top of the gels and the cells were left to acclimate to the gels overnight. The medium on top of the gel in each well was then replaced with 0.25 mL of medium containing nanoparticles (final total concentration ∼10^11^ NPs/mL). The cells were then incubated with the nanoparticles for 4 hours after which the media was replaced with 0.25 mg/mL liberase TL and incubated for 25 minutes. The gels with the liberase TL were then moved to microcentrifuge tubes and incubated for a further 5 minutes on a rotating shaker until the gels were fully dissolved. The cells were then collected via centrifugation after which they were stained and fixed in the same manner as used for cells in 2D culture.

### Statistical Analysis

GraphPad Prism 9 (GraphPad Software) was used for graphing and statistical analysis of the data. A one-way analysis of variance (ANOVA) with a Tukey post hoc correction was performed for analysis between groups. For groups with non-Gaussian distributions, Kruskal-Wallis with Dunn’s correction was used. Specific statistical tests used for each data set are noted in the figure caption. Bar graphs show mean and standard deviation and box and whiskers plots show median value and 5th percentile up to 95th percentile. Statistical significance was assessed at p < 0.05.

## Results

### Generation of tissue-specific ECM hydrogels

As noted, previous studies by our group and others have examined the interactions of nanoparticles with ECM using models such as collagen and Matrigel that are not wholly representative of the native matrix. Purified collagen scaffolds lack many important ECM components such as fibronectin and proteoglycans while Matrigel, as a basement membrane model, is not representative of the interstitial matrix.^24–27^ There are also distinct differences in ECM composition and organization between different types of tissue that are not represented in these models.^28^ To generate a model ECM with tissue-specific properties, we prepared dECM hydrogels derived from porcine lung, liver, and SIS for use in our studies. A series of chemical processes combined with mechanical agitation were used to obtain decellularized tissue from the porcine lung and liver (**Fig. 1A**). To obtain decellularized SIS, we scraped the cells from the submucosa and sterilized the scaffold using PAA (**Fig. 1B**). We validated the successful decellularization of the tissues by quantifying the dsDNA content. Decellularized tissue contained dsDNA concentrations significantly lower than native tissue (<100 ng/mg and >6000 ng/mg respectively). We next characterized the biochemical makeup of the decellularized tissue by determining their collagen and sulfated GAG (sGAG) content. The soluble collagen content was similar among all types of decellularized tissue with around 50 percent of the tissue made up of soluble collagen (**Fig. 1C**). Decellularized small intestine contained the highest concentration of sGAG which was more than twice the amount of either decellularized liver or lung (**Fig. 1D**).

**Figure 1.**
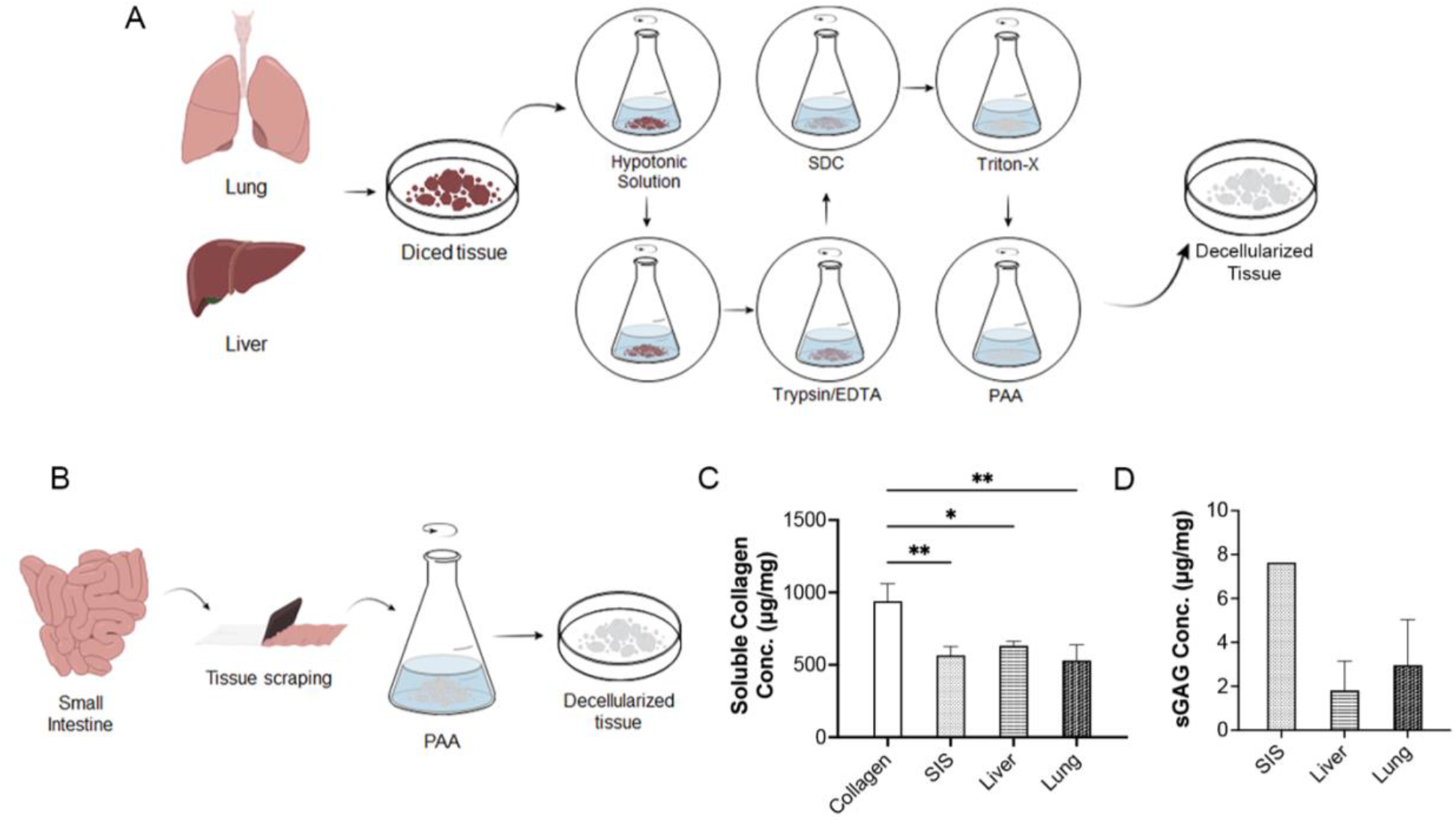
Decellularization and biochemical characterization of lung, liver, and small intestine tissue. Method for the chemical and mechanical decellularization of (**A**) lung, liver, and (**B**) small intestine tissue. Soluble collagen content of decellularized tissue prior to gelation compared to commercial collagen. Concentration of sulfated GAG in different types of tissue normalized to the amount of tissue. Data sets in (**C, D**) statistically analyzed with one-way analysis of variance (ANOVA) and a Tukey post hoc correction: *p < 0.05, **p < 0.01, and not significant unless otherwise noted.

### Viscoelastic properties of dECM hydrogels

During cancer progression, ECM within in the tumor microenvironment increases in overall density and stiffness due to an upregulation of collagen and other ECM components.^2^ To mimic this increase in ECM viscoelasticity, we created hydrogels with varying ECM content and analyzed their viscoelastic properties using both bulk- and micro-rheological measurements. Bulk measurements showed that the elastic modulus (G’), a measure of gel stiffness, of the ECM hydrogels increased with increasing ECM content as expected (**Fig. 2A**). While the liver dECM hydrogels generally had the highest G’, differences were not statistically significant. To analyze any micro-scale differences between the hydrogels, we used particle tracking microrheology to characterize the biophysical properties of dECM hydrogels at varying concentrations (**Fig. 2B,C**). Based on our microrheological analysis, G’ significantly increased with increasing dECM concentrations of 8 mg/mL (**Fig. 2B**). There were also notable differences in measured G’ between dECM derived from different tissue type (G’_SIS_ > G’_lung_; G’_liver_ > G’_lung_). Based on G’ determined via microrheology, we estimated the pore sizes (ξ) in dECM hydrogels which ranged from 500– 1500 nm (**Fig. 2C**). We found significant differences in pore sizes between tissue types (ξ_SIS_ < ξ_lung;_ ξ_liver_ < ξ_lung;_). In addition, the pore size significantly decreased with increasing dECM concentration.

**Figure 2.**
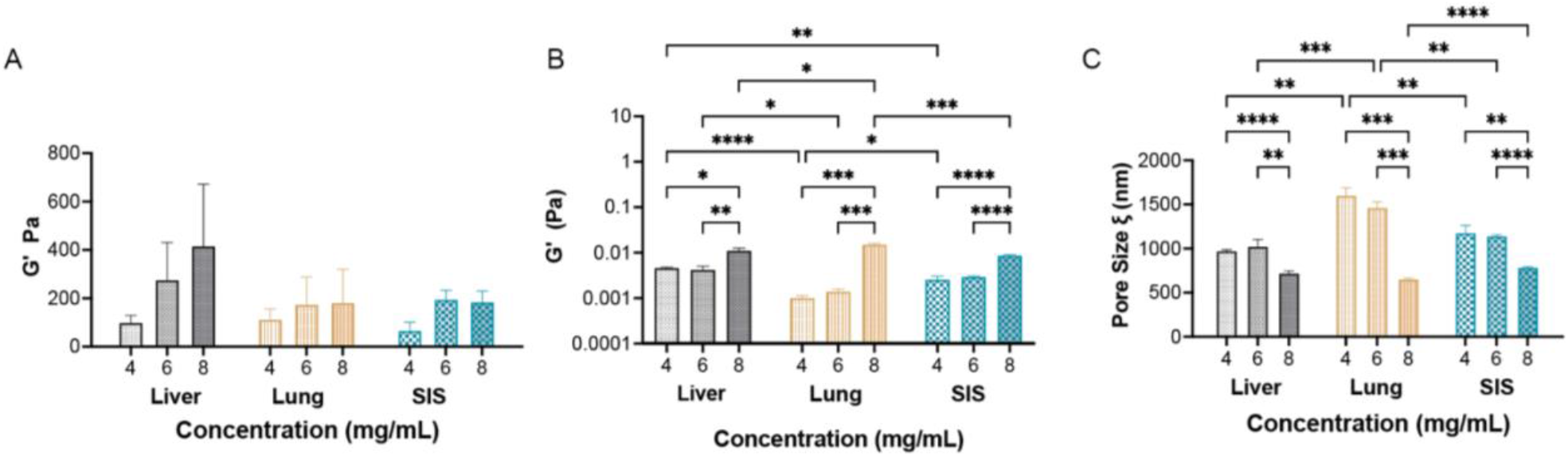
Viscoelastic properties of ECM hydrogels. (**A**) Elastic moduli for 4, 6, and 8 mg/mL SIS, liver, and lung hydrogels obtained using a bulk rheometer with parallel plate geometry. (**B**) Elastic moduli for 4, 6, and 8 mg/mL SIS, liver, and lung hydrogels as calculated via microrheology. (**C**) Estimated pore size for 4, 6, and 8 mg/mL SIS, liver, and lung gels from microrheological measurements. Data sets in (**A-C**) statistically analyzed with one-way analysis of variance (ANOVA) and Dunnett’s test: *p < 0.05, **p < 0.01, ***p < 0.001, ****p<0.0001 and not significant unless otherwise noted.

### Nanoparticle diffusion in ECM

We analyzed the effects of ECM source and PEGylation type on NP transport using fluorescence video microscopy and multiple particle tracking analysis to measure the diffusion of NPs with or without PEG coatings in varying concentrations of lung, liver, and SIS dECM hydrogels or in collagen gels (**Fig. 3**). Collagen gel concentrations were matched to the approximated collagen content in the dECM hydrogels. Nanoparticles were coated with linear 2 kDa PEG (PEG-L2), linear 5 kDa PEG (PEG-L5), linear 10 kDa (PEG-L10), and branched 10 kDa PEG (PEG-B10). Qualitatively, non-PEGylated NPs appeared immobile in all gel types while significant mobility was noted for the densely PEGylated NPs (**Fig. 3A**). While the non-PEGylated NPs had little to no mobility in all hydrogel types (**Fig. 3B**), we found the diffusion rate, as measured by MSD, of PEG-L5 NPs was higher in dECM hydrogels as compared to collagen alone (**Fig. 3C,D**). When comparing PEGylation types in the highest concentration of liver hydrogels, we found PEG-L10 and PEG-B10 NPs achieved significantly higher diffusion rate compared to unPEGylated and PEG-L2 NPs (**Fig. 3E**). Increasing the concentration of liver-derived dECM hydrogels significantly impeded PEGylated NP mobility as expected (**Fig. 3F**). To enable comparison across PEG and tissue types, we also calculated a normalized MSD_1s_ with respect to UnPEGylated NPs, which had little to no movement in all hydrogels regardless of tissue type or concentration, and thus indicates the fold increase in mobility for each NP (**Fig. 3G-I**). While all PEGylated NPs had significantly higher diffusion in all hydrogels compared to the non-PEGylated NPs, the type of PEG that performed best depended on the type of tissue and concentration of ECM used to form the hydrogels. Overall, we find high-density PEGylation with lower molecular weight PEG (PEG-L2 or PEG-L5) offers significant enhancement in NP transport through different ECM types. However, exceptions to this trend were observed for instance at the highest concentration hydrogels, where the PEG-B10 NPs diffused the fastest in the SIS hydrogels (**Fig. 3I**). In addition to these summary plots, we also provide measured MSD for NPs diffusing in dECM and collagen across all experimental conditions tested in the supplement (**Figs. S1-3**).

**Figure 3.**
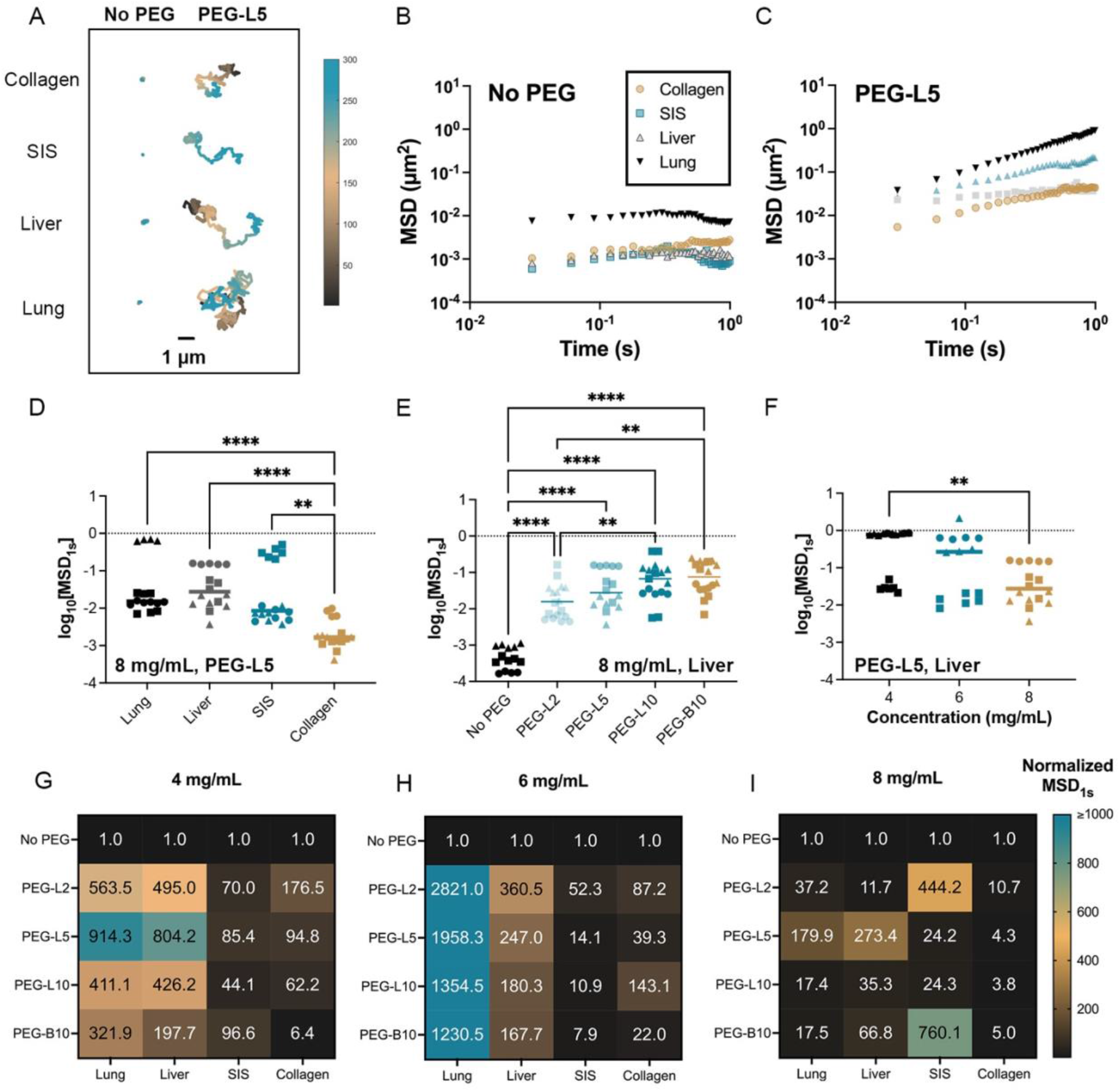
Nanoparticle diffusion in interstitial ECM models. (**A**) Representative trajectories of non-PEGylated (no PEG) and linear 5 kDa PEGylated (PEG-L5) NPs in ECM hydrogels. Traces show 10 seconds of motion with a color scale to indicate time. The scale bar represents 1 µm. Average MSD over time for (**B**) non-PEGylated and (**C**) PEG-L5 NPs in 8 mg/mL lung dECM, liver dECM, SIS dECM, and collagen hydrogels. (**D**) Diffusion of PEG-L5 NPs represented as the median log based 10 of MSD at *τ* = 1 s (log_10_[MSD_1s_]) in lung dECM, liver dECM, SIS dECM, and collagen hydrogels. (**E**) Diffusion of uncoated NP, PEG-L5 NP, and NPs coated with 2 kDa linear PEG (PEG-L2), 10 kDa linear PEG (PEG-L10), and 10 kDa branched PEG (PEG-L10) in 8 mg/mL liver dECM hydrogels. (**F**) Diffusion of PEG-L5 NPs in liver dECM hydrogels with varied concentration from 4–8 mg/mL. For (**D-F**), data sets statistically analyzed with one-way analysis of variance (ANOVA) and Dunnett’s test: *p < 0.05, **p < 0.01, ***p < 0.001, ****p<0.0001. **(G-I)** Summary heatmap of MSD_1s_ for each NP type normalized to MSD_1s_ for unPEGylated NP (no PEG) in dECM and collagen hydrogels at (**G)** 4 mg/mL, (**H)** 6 mg/mL, and (**I)** 8 mg/mL concentrations. At least 3 videos in different regions of the gel were acquired and analyzed in each experiment. An average of >160 particles were tracked per sample tested.

### Nanoparticle uptake in cell embedded ECM hydrogels

ECM-rich stromal regions within tumors and healthy tissues can impact NP diffusion.^42,43^ However, it is currently unclear in what manner and to what extent these regions affect NP transport and whether these differences are tissue dependent. To simulate the *in vivo* environment where stromal cells would be dispersed within interstitial ECM, we embedded human cancer cells within ECM hydrogels derived from different tissues (**Fig. 4A**). We administered NPs to the cell-embedded hydrogels and measured the percentage of cells that had bound or internalized NPs. The percent uptake was normalized to the uptake of non-PEGylated NPs to facilitate comparison between tissue types. For cells not embedded in ECM hydrogels which were cultured on plastic in a 2D environment, virtually all cells were able to uptake NPs regardless of the type of NP administered (**Fig. 4B**). However, the total amount of NPs taken up by the cells was affected by the type of PEG coated on the NP surface as observed through their geometric mean fluorescence intensity (gMFI). The PEG-L2 NPs had a significantly higher gMFI compared to all other types of NPs, but there were not any statistical differences between the remaining NP types (**Fig. 4C**). In liver hydrogels, the non-PEGylated NPs were uptaken by a significantly lower percent of cells compared to the PEGylated NPs while there were no significant differences between the uptake of the PEGylated NPs (**Fig. 4D**). The PEGylated NPs also had a significantly higher uptake in cells cultured in lung hydrogels compared to the non-PEGylated NPs except the PEG-L10 NPs (**Fig. 4E**). Additionally, PEG-L2 NPs had significantly higher cellular uptake in lung dECM hydrogels compared to all other types of PEGylated NPs. Both PEG-L2 and PEG-B10 NPs had a significantly higher uptake than the PEG-L5 or PEG-L10 NPs in cells embedded in SIS dECM (**Fig. 4F**). The relative number of NPs taken up by the cells as measured by the gMFI showed a noticeable shift in the peaks for the PEGylated NPs compared to the non-PEGylated NPs (**Fig. 4G-I**). However overall, the gMFI of cells in dECM hydrogels followed similar trends as the overall percent NP uptake for each PEG type (**Fig 4K-L**).

**Figure 4.**
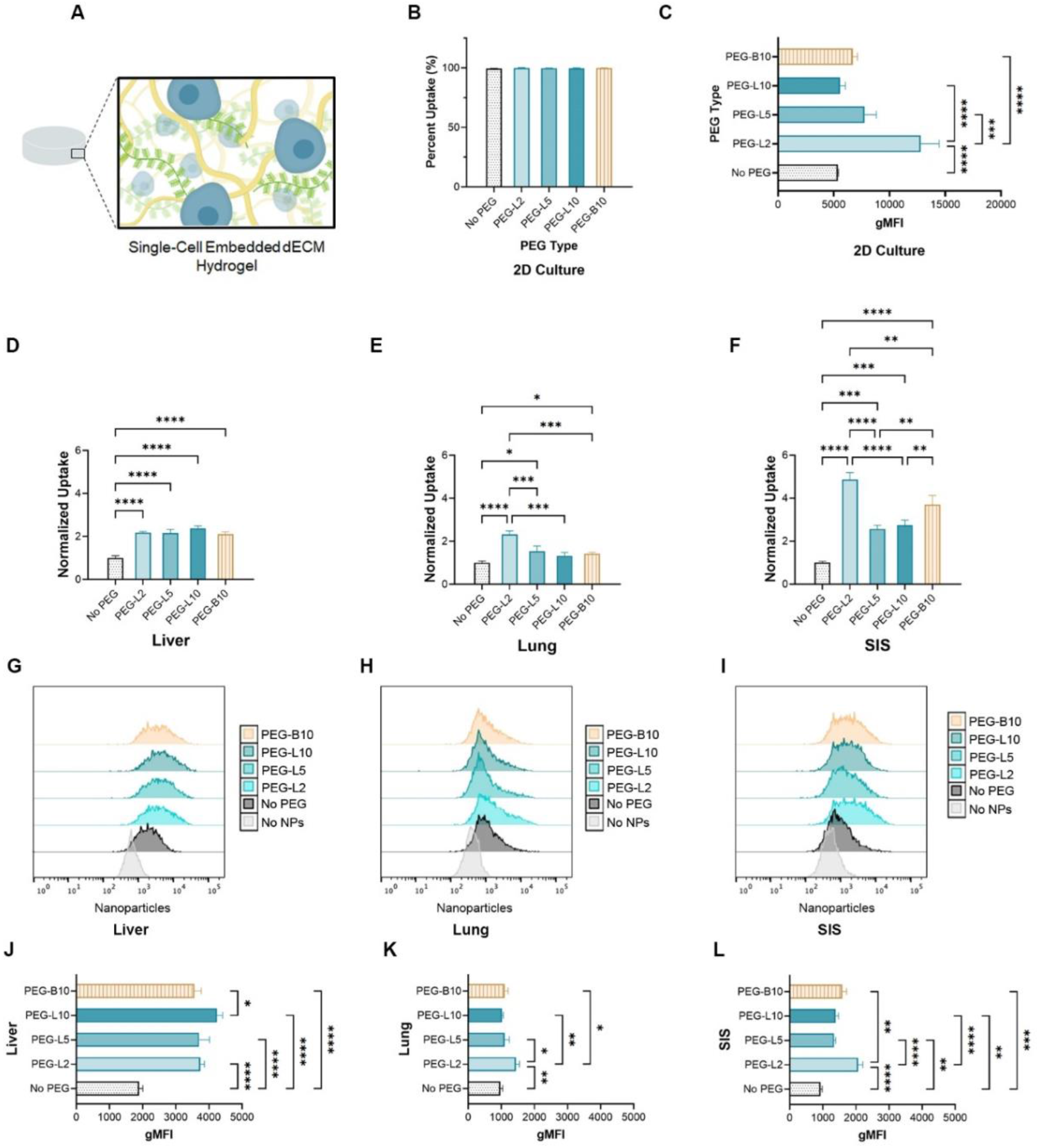
NP uptake in cell-embedded dECM hydrogels derived from different tissues. (**A**) Schematic of an ECM hydrogel with cells interspersed throughout the network. (**B**) Percent of cells cultured on plastic in a 2D environment with uncoated NPs (no PEG) and NPs coated with 2 kDa linear PEG (PEG-L2), 5 kDa linear PEG (PEG-L5), 10 kDa linear PEG (PEG-L10), and 10 kDa branched PEG (PEG-L10). (**C**) Geometric mean fluorescence intensity (gMFI) of NPs in cells cultured on plastic in a 2D environment. (**D-F**) Percent of cells with NPs normalized to the result for non-PEGylated NPs in 6 mg/mL liver, lung, and SIS hydrogels respectively. (**G-I**) Fluorescence spectra of nanoparticles in 6 mg/mL liver, lung, and SIS hydrogels respectively. (**J-L**) Geometric mean fluorescence intensity of nanoparticles in cells embedded in 6 mg/mL liver, lung, and SIS hydrogels respectively. Data sets statistically analyzed with one-way analysis of variance (ANOVA) and a Tukey post hoc correction: *p < 0.05, **p < 0.01, ***p < 0.001, ****p<0.0001 and not significant unless otherwise noted.

We varied the concentration of dECM hydrogels to determine whether and to what extent there was a concentration dependence to the NP-ECM interactions. In **Fig. 5**, we show the outcomes of our studies in liver dECM hydrogels with varied concentration. Additional results for cellular uptake in lung and SIS dECM hydrogels at low (4 mg/mL) and high (8 mg/mL) concentrations are provided in the supplement (**Fig. S4, S5**). The overall uptake had a similar trend between NP types regardless of the concentration of liver dECM (**Fig. 5A-C)**. The impact of PEGylation strategy was most evident in the 4mg/mL ECM gels compared to the higher concentration gels, with a significantly higher uptake compared to non-PEGylated NPs (**Fig. 5A**). A noticeable shift was observed for the NP fluorescence of PEGylated NPs compared to non-PEGylated NPs for all concentrations of the ECM hydrogels (**Fig. 5D-F**). For the 4 mg/mL ECM hydrogels, which were the lowest concentration measured, there was also a notably greater shift in the spectra of the PEG-L2 and PEG-B10 NPs compared to other NP types (**Fig. 5D**). Based on measured gMFI, significantly greater numbers of PEGylated NPs were uptaken in dECM-embedded cells compared to non-PEGylated NPs (**Fig. 5G-I)**. At the highest dECM concentration (8 mg/mL), there was notably lower gMFI for both PEGylated and unPEGylated NP, suggesting a lower number of NPs taken up per cell (**Fig. 5G-I**).

**Figure 5.**
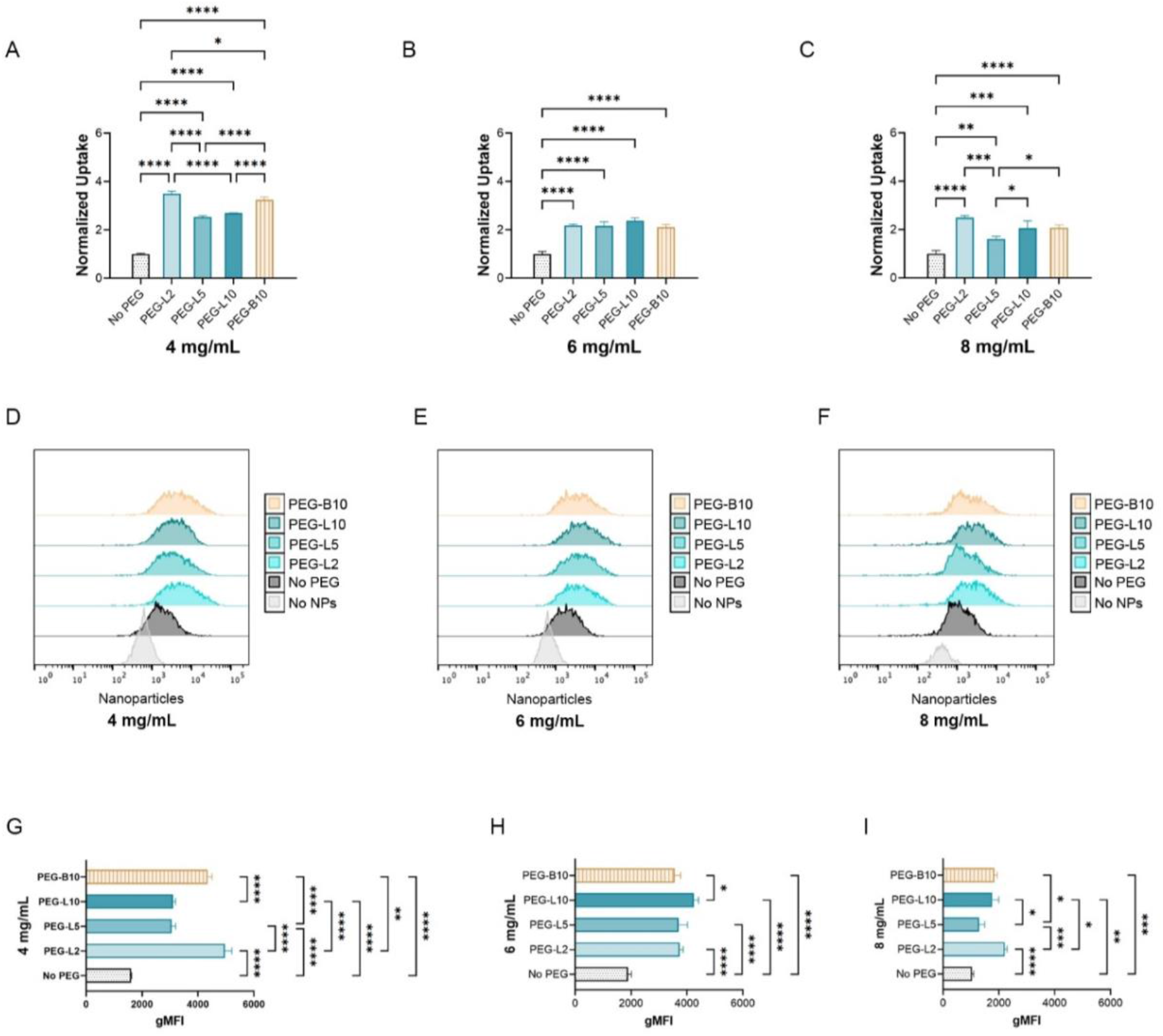
NP uptake in cell-embedded liver dECM hydrogels of varying concentrations. (A-C) Percent of cells with NPs that are uncoated (no PEG) or coated with 2 kDa linear PEG (PEG-L2), 5 kDa linear PEG (PEG-L5), 10 kDa linear PEG (PEG-L10), and 10 kDa branched PEG (PEG-L10). Results normalized to non-PEGylated NPs in 4, 6, and 8 mg/mL liver dECM hydrogels respectively. (D-F) Fluorescence spectra of nanoparticles in 4, 6, and 8 mg/mL liver hydrogels resepctively. (G-I) Geometric mean fluorescence intensity of nanoparticles in cells embedded in 4, 6, and 8 mg/mL liver hydrogels respectively. Data sets statistically analyzed with one-way analysis of variance (ANOVA) and a Tukey post hoc correction: *p < 0.05, **p < 0.01, ***p < 0.001, ****p<0.0001 and not significant unless otherwise noted.

## Discussion

In this study, dECM hydrogels were used to model tissue-specific ECM composition and density to determine how these differences impact NP transport following different PEGylation strategies. The overall structure, organization, and composition of ECM can vary greatly depending on the tissue from which it is derived.^28^ We hypothesized that these differences would significantly impact NP transport and would be related to the relative abundance of ECM components (e.g. collagen, sGAG) as well as their biophysical properties. We found dECM hydrogels from different tissues possessed similar viscoelasticity and pore size of the networks presumably due to the similar amounts of fibrillar collagen in each scaffold. We also found that PEGylated NP diffusion in collagen gels was significantly more hindered in comparison to dECM gels. This suggests the collagen matrix poses a greater barrier to NP diffusion in the absence of non-collagen components. Collagen is known to interact with other proteoglycans in the ECM which can alter collagen fibril assembly and organization.^44,45^ Thus, specific and non-specific interactions between collagen and other ECM components may interfere with NP association to collagen fibers. However, additional research is warranted into the types of non-collagen species present and their molecular features to provide additional insights into these interactions.

The tissue-specific origin of dECM hydrogels affected composition and self-assembly, which in turn may have influenced NP diffusion. When comparing between tissues, the lung dECM hydrogels had, on average, a larger pore size than all other hydrogels based on our microrheological assessment, which may have contributed to the enhanced mobility of NPs in these gels compared to liver and SIS dECM hydrogels. SIS dECM had the greatest sGAG content and generally the lowest NP mobility among the different dECM hydrogels tested. It is possible that the higher sGAG content in the SIS dECM gels resulted in greater interstitial crowding,^46,47^ creating more steric hindrance for the NPs and obstructing their diffusion through the network. However, we recognize these differences could also, at least in part, be attributed to the method of preparation for SIS dECM as compared to lung and liver dECM. As previously noted, further work is required to determine which individual or combinations of non-collagen components play the most substantial role in modulating NP penetration through the ECM. Given the hundreds of proteins present in ECM,^48^ this is beyond the scope of our current study and will be pursued in future studies. Despite these limitations, these studies establish tissue-specific ECM organization and composition can markedly impact NP diffusion.

Previous studies have shown that PEG molecular weight can significantly impact the mobility of NPs in ECM and other hydrogel networks.^7,23,49^ We also found in our previous work that a high density coating of branched PEG significantly improves NP diffusion in Matrigel, a model of the basement membrane portion of the extracellular matrix.^23^ We hypothesized that this dense coating of branched PEG could offer the same advantages in the interstitial portions of the ECM due to its enhanced shielding effects. In our models of the interstitial matrix, we found that the ideal PEG coating depended on the tissue source and ECM concentration, but linear PEG with a molecular weight of 5 kDa or lower tended to perform better in most types of ECM and collagen. Interestingly, the branched PEG NPs, proved to be the exception to this observation within SIS hydrogels and tended to perform better than all other types of NPs in the SIS gels while they had a comparable diffusion profile to PEG-L10 NPs in lung and liver hydrogels. It is possible that the branched PEG offered additional protection of the NPs from interactions with the sGAGs that were present at higher concentrations in the SIS hydrogels compared to the lung and liver hydrogels.

During disease progression, there is an upregulation of ECM components, including collagen and HA.^17^ Therefore, we varied the concentration of ECM in our hydrogels to model this increase which can cause increased ECM viscoelasticity and reduce NP mobility. As expected, NPs had decreased mobility with increasing concentrations of ECM. This can be largely attributed to a reduction in pore size and increase in stiffness of the hydrogel networks with increasing ECM concentration as evidenced by our microrheological assessments which showed a decreased pore size and increased elastic modulus for higher ECM concentrations. Interestingly, mobility of the NPs was not greatly impacted by the increasing ECM concentration for lung and liver dECM hydrogels until they encountered the highest concentration gels while the NPs exhibited significantly reduced diffusion in the intermediate concentration 6 mg/mL SIS dECM gels compared to the lowest concentration 4 mg/mL SIS dECM gels. The greater impact of increasing concentration in SIS gels compared to lung and liver gels may be attributed to the greater sGAG content present in SIS which, as mentioned earlier, may increase steric hindrance, reducing NP diffusion.

Many studies have shown that PEGylating NPs reduces cell uptake which can reduce efficacy of drug carriers.^22,50^ However, traditional *in vitro* studies in a 2D setting do not account for the role that the ECM microenvironment may play in the ability of NPs to reach target cells. The role of tissue-specific ECM barrier properties on NP drug delivery is also difficult to directly assess in more complex *in vivo* animal models. To determine the role the ECM and tissue source plays in NPs reaching target cells, we created a 3D cell culture model using dECM hydrogels with cells embedded throughout the network. We hypothesized that in the presence of ECM, PEGylated NPs would have a higher uptake in cells compared to non-PEGylated NPs. We used a high concentration of NPs which enabled virtually all cells in a 2D environment to uptake NPs to determine to what extent the NPs were impacted by the ECM in our 3D model. As we had hypothesized, all PEGylated NPs had greater uptake than the non-PEGylated NPs. We also found reduced NP uptake in cells embedded within ECM gels of increasing concentration. The NP uptake generally followed a similar trend to our MPT results with lower molecular weight PEG performing best. The shorter PEG chains on the PEG-L2 NPs may provide an optimal balance between effectively transporting through the different types of ECM while simultaneously enabling greater uptake upon reaching the target cells. However, we also observed a tissue-specific enhancement of cellular uptake for PEG-B10 NPs in SIS dECM gels which may be related to enhanced ECM penetration in this tissue type. Furthermore, these data suggest that ECM likely influences NP delivery efficiency by not only acting as a physical barrier to penetration but also by affecting cell uptake competency. While we have shown that PEGylation can increase overall NP uptake in target cells in the presence of ECM, additional studies into the overall distribution of NP uptake in cells embedded in dECM gels would be useful in determining how deeply the NPs penetrated and interact with cells within the 3D matrix.

## Conclusion

The results presented here show that NP transport is significantly impacted by tissue-specific properties of ECM. We also confirmed that PEGylation improves uptake in target cells in the presence of ECM. We demonstrated that low molecular weight PEG coatings generally enables NP transport and cellular uptake within ECM across the tissues tested. However, tissue-specific ECM barrier functions were also noted where alternative PEGylation strategies may improve NP delivery to target cells. Future work in other ECM-enriched tissues (e.g. brain ECM) and in the context of disease (e.g. tumor-derived ECM) may provide further insight on optimal formulation strategies to overcome the ECM as a barrier to effective NP drug delivery. Taken together, this work provides a framework to evaluate the impacts of the ECM on nanoparticle delivery systems with varied size, shape, surface chemistry, and cellular targeting strategies which can aid in their preclinical development.

## Acknowledgments

This project was funded by the UMD-NCI Partnership for Integrative Cancer Research and the National Science Foundation (CAREER Award 2047794). We acknowledge the BioWorkshop core facility in the Fischell Department of Bioengineering at the University of Maryland for use of their dynamic light scattering instrument and flow cytometer.

