## Supporting Information for "The Extracellular Matrix Limits Nanoparticle Diffusion and Cellular Uptake in a Tissue-Specific Manner"

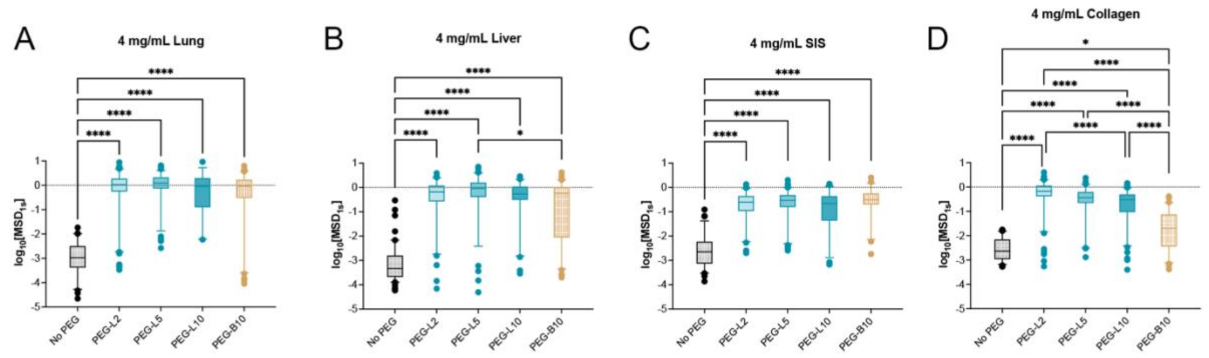


**Figure S1. Nanoparticle diffusion in 4 mg/ml dECM and collagen hydrogels.** Measured log based 10 of mean squared displacement at time scale of 1 second (log10[$MSD_{1s}$]) for (**A**) NP diffusion in 4 mg/mL liver hydrogels. (**B**) NP diffusion in 4 mg/mL lung hydrogels. **(C)** NP diffusion in 4 mg/mL SIS hydrogels. (**D)** NP diffusion in 4 mg/mL collagen hydrogels. ****p<0.0001 and *p<0.05 by Kruskal-Wallis with Dunn's correction. At least 3 videos in different regions of the gel were acquired and analyzed in each experiment. An average of >160 particles were tracked per sample tested.

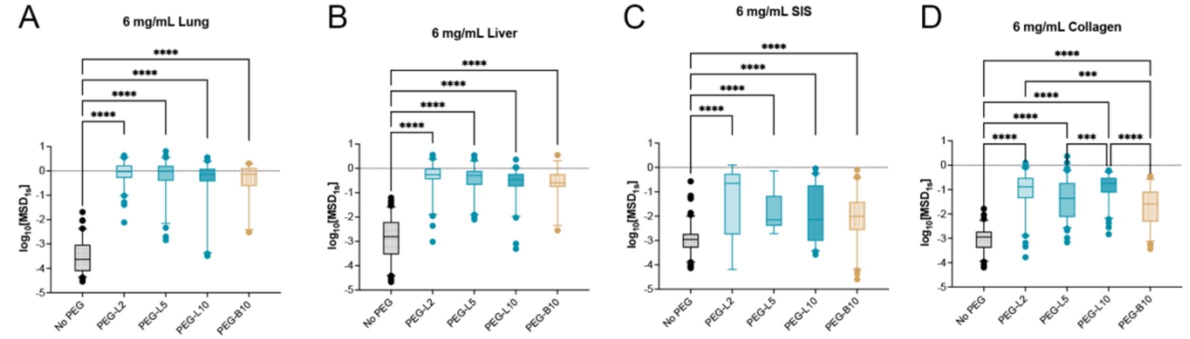


**Figure S2. Nanoparticle diffusion in 6 mg/ml dECM and collagen hydrogels.** Measured log based 10 of mean squared displacement at time scale of 1 second (log10[$MSD_{1s}$]) for (**A**) NP diffusion in 6 mg/mL liver hydrogels. (**B**) NP diffusion in 6 mg/mL lung hydrogels. **(C)** NP diffusion in 6 mg/mL SIS hydrogels. (**D)** NP diffusion in 6 mg/mL collagen hydrogels. ****p<0.0001 and ***p<0.001 by Kruskal-Wallis with Dunn's correction. At least 3 videos in different regions of the gel were acquired and analyzed in each experiment. An average of >160 particles were tracked per sample tested.

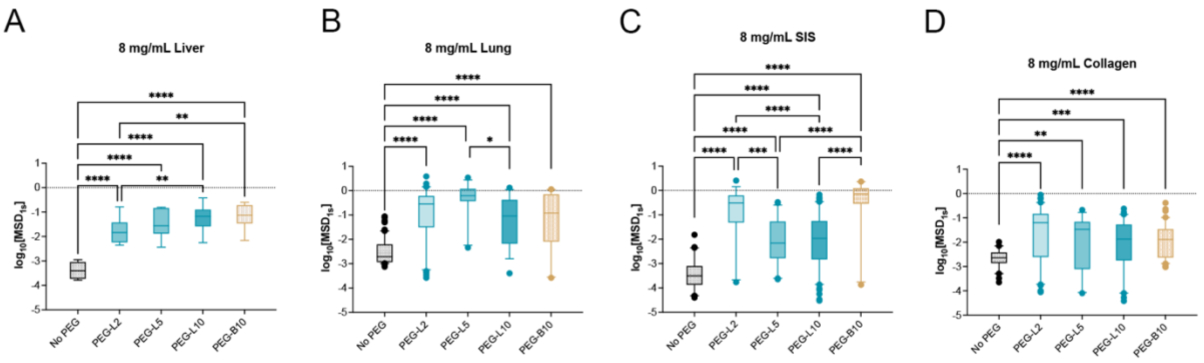


**Figure S3. Nanoparticle diffusion in 8 mg/ml dECM and collagen hydrogels.** Measured log based 10 of mean squared displacement at time scale of 1 second (log10[$MSD_{1s}$]) for (**A**) NP diffusion in 8 mg/mL liver hydrogels. (**B**) NP diffusion in 8 mg/mL lung hydrogels. **(C)** NP diffusion in 8 mg/mL SIS hydrogels. (**D)** NP diffusion in 8 mg/mL collagen hydrogels. ****p<0.0001, ***p<0.001, **p<0.01, and *p<0.05 by Kruskal-Wallis with Dunn's correction. At least 3 videos in different regions of the gel were acquired and analyzed in each experiment. An average of >160 particles were tracked per sample tested.

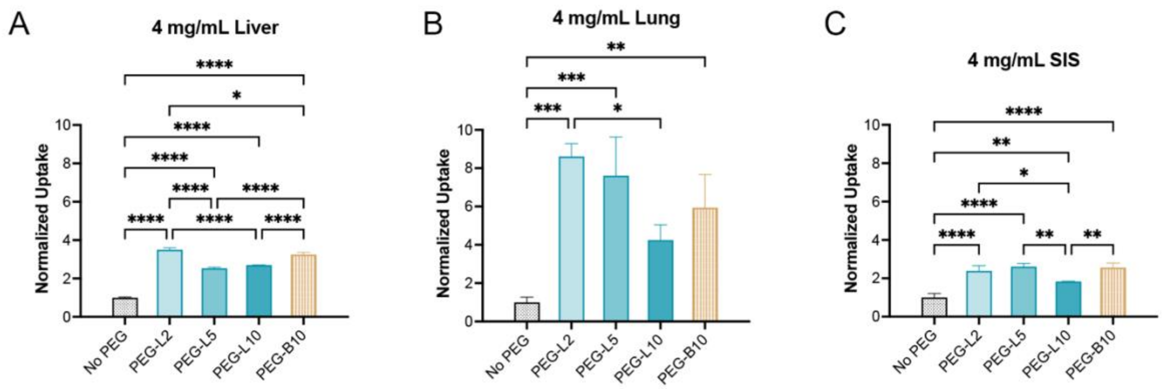


**Figure S4. NP uptake in cell-embedded 4 mg/mL dECM hydrogels derived from different tissues.** (A-C) Percent of cells with NPs normalized to the result for non-PEGylated NPs in 4 mg/mL liver, lung, and SIS hydrogels respectively. Data sets statistically analyzed with one-way analysis of variance (ANOVA) and a Tukey post hoc correction: ns = not significant; *p < 0.05; **p < 0.01; ***p < 0.001; ****p < 0.0001.

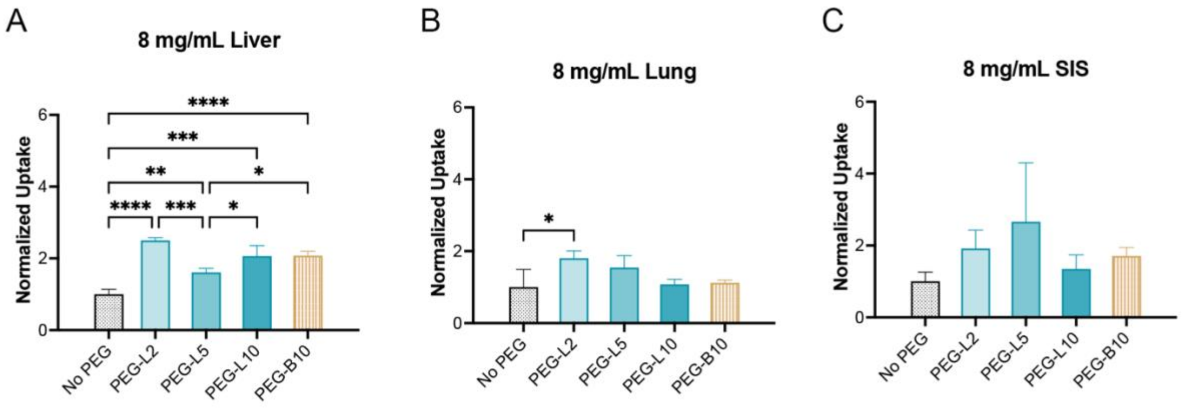


**Figure S5. NP uptake in cell-embedded 8 mg/mL dECM hydrogels derived from different tissues.** (A-C) Percent of cells with NPs normalized to the result for non-PEGylated NPs in 8 mg/mL liver, lung, and SIS hydrogels respectively. Data sets statistically analyzed with one-way analysis of variance (ANOVA) and a Tukey post hoc correction: ns = not significant; *$p < 0.05$; **$p < 0.01$; ***$p < 0.001$; ****$p < 0.0001$.
